# Artificial Neural Networks for classification of single cell gene expression

**DOI:** 10.1101/2021.07.29.454293

**Authors:** Jiahui Zhong, Minjie Lyu, Huan Jin, Zhiwei Cao, Lou T. Chitkushev, Guanglan Zhang, Derin B. Keskin, Vladimir Brusic

**Affiliations:** School of Computer Science, University of Nottingham Ningbo China; School of Life Sciences and Technology, Tongji University, Shanghai, China; HiLab, Metropolitan College, Boston University, Boston, USA; Translational Immuno-Genomics Lab, Dana-Farber Cancer Institute, Harvard Medical School, Boston, USA

**Author notes:** Correspondence, School of Computer Science, University of Nottingham Ningbo China, Ningbo 315100, China.

**Keywords:** Cell type classification, ANN training, healthy PBMC samples, incremental learning, SCT data, PBMC cell types

## Abstract

**Background:** Single-cell transcriptome (SCT) sequencing technology has reached the level of high-throughput technology where gene expression can be measured concurrently from large numbers of cells. The results of gene expression studies are highly reproducible when strict protocols and standard operating procedures (SOP) are followed. However, differences in sample processing conditions result in significant changes in gene expression profiles making direct comparison of different studies difficult. Unsupervised machine learning (ML) uses clustering algorithms combined with semi-automated cell labeling and manual annotation of individual cells. They do not scale up well and a workflow used on a specific dataset will not perform well with other studies. Supervised ML classification shows superior classification accuracy and generalization properties as compared to unsupervised ML methods. We describe a supervised ML method that deploys artificial neural networks (ANN), for 5-class classification of healthy peripheral blood mononuclear cells (PBMC) from multiple diverse studies.

**Results:** We used 58 data sets to train ANN incrementally – over ten cycles of training and testing. The sample processing involved four protocols: separation of PBMC, separation of PBMC + enrichment (by negative selection), separation of PBMC + FACS, and separation of PBMC + MACS. The training data set included between 85 and 110 thousand cells, and the test set had approximately 13 thousand cells. Training and testing were done with various combinations of data sets from four principal data sources. The overall accuracy of classification on independent data sets reached 5-class classification accuracy of 94%. Classification accuracy for B cells, monocytes, and T cells exceeded 95%. Classification accuracy of natural killer (NK) cells was 75% because of the similarity between NK cells and T cell subsets. The accuracy of dendritic cells (DC) was low due to very low numbers of DC in the training sets.

**Conclusions:** The incremental learning ANN model can accurately classify the main types of PBMC. With the inclusion of more DC and resolving ambiguities between T cell and NK cell gene expression profiles, we will enable high accuracy supervised ML classification of PBMC. We assembled a reference data set for healthy PBMC and demonstrated a proof-of-concept for supervised ANN method in classification of previously unseen SCT data. The classification shows high accuracy, that is consistent across different studies and sample processing methods.

## Background

Classification of single cells is an essential step for analyzing the composition of tissues and better understanding of the cellular basis of health and disease. Accurate classification of cell types and subtypes, along with the identification of their gene and protein expression patterns, enable understanding the nature and the extent of biological processes in the observed tissues [1]. This knowledge allows for medical applications of single cell technologies: diagnostic and prognostic applications, and disease treatment selection [2]. Peripheral blood mononuclear cells (PBMC) are circulating immune cells with a single round nucleus in the blood and are common diagnostic and prognostic targets [3]. PBMC are important targets of single-cell studies because they are indicators of immune status and are studied in cancer [4–5], infectious disease [6–7], and autoimmunity [8–10], among others. Five major cell types that constitute PBMC are B cells (BC), dendritic cells (DC), monocytes (MC), natural killer cells (NK), and T cells (TC) [11]. Gene expression profiles in PBMC that circulate in blood were shown to be different from the tissue resident PBMC [12]. This suggests that gene expression differences can also be used to identify the tissue of origin of resident PBMC [13].

Normal ranges (reference values) of the numbers of specific cell types or subtypes in PBMC vary by 5 to 20 folds in healthy individuals [14]. Their transcriptome profiles show high variation, primarily resulting from sample processing steps [15] and the health/disease status of the tissue [3,16–17]. More than 120 cell subtypes of PBMC have been described [18], but current descriptions of PBMC subsets are incomplete and the efforts to define them are ongoing [19–20]. In addition to the inherent biological differences, each step in the process of peripheral blood sampling, storage, preparation, and measurement as well as their duration will change gene expression in single cells [21–23]. At present, uniform and strict standards have been established for sample collection, preparation, and storage of PBMC [21,24]. Standard operating procedures (SOP) have been defined and established for the latest single cell transcriptomics (SCT) technologies [25], providing for improved reproducibility of SCT studies. The combination of advanced SCT technologies and the rapidly increasing availability of data sets provide a basis for defining cell types and subtypes by SCT gene expression profiles from diverse datasets.

The first report of single cell gene expression was published in 2009 [26]. Major breakthroughs in microfluidics and cell labeling methods have enabled high-throughput of single cells, rapid standardized SCT gene expression measurement, and analysis [27–29]. The conventional classification rules applied to cell populations are mainly qualitative and are based on lineage, phenotypic markers, and simple, functional properties [30]. The SCT uses gene expression and quantitative methods to define cell types and precisely describe their lineage, phenotype, function, and various states [30]. Such cellular gene expression profiles and their variants (due to different sample processing methods) are cataloged in single cell atlases [31–32]. Bulk-sequencing methods produce mean gene expression values of millions of cells. In contrast, SCT produces gene expression profiles characteristic of cell sets defined by a much finer grouping of cells that share origin, function, subtype, and biological status [33].

The scRNA-seq data produced using 10x Genomics platform is the current method of choice because it combines high throughput (up to 40,000 cells in a single experiment), affordable cost, and rapid turnaround (1-2 days from sample collection to results) [34]. When the cell viability is greater than 90%, the cell capture rate of one single sample can reach 65% (10x protocol). The 10x SCT data is represented by a high-dimensional sparse matrix. A single cleaned 10x SCT data set (sparse matrix) may have 10^9^–10^10^ data points because it has up to 10^5^ columns representing individual cells and >30,000 rows representing features (gene counts). We have observed that 90-99% of the values are zero [12]. Since 2017, with the emergence of the 10x Genomics platform, the large-scale unified 10x scRNA-seq data sets have been generated and have grown exponentially with more than 14,000 10x data sets available in GEO data repository (www.ncbi.nlm.nih.gov/geo), as of May 2021.

Currently, the analysis of SCT data focuses on single cell annotation and classification aimed at understanding biological mechanisms, such as cellular differentiation, tissue distribution of cells, the discovery of new biomarkers, detection of rare cell types, assessment of tumor heterogeneity, detecting gene activation pathways related to pathology, and detecting molecular and cellular responses to therapeutic interventions [35–37]. The characteristics of SCT data – large size, sparseness, sensitivity to sample processing and experimental conditions, biases and random errors in data, and lack of reference data sets – require advanced statistical and machine learning (ML) techniques essential for the analysis of sparse matrices (downstream analysis). SCT data sets are produced using various sample processing conditions and they represent many different biological states, making SCT data highly heterogeneous. The lack of reference data sets mandates the use of unsupervised ML approaches, predominantly unsupervised clustering [37]. Unsupervised ML methods are broadly used for labeling and classification of single cells either alone [38] or in combination with supervised ML methods [39]. Unsupervised ML methods deploy a combination of clustering algorithms with semi-automated labeling and manual annotation of individual cells. They do not scale up well, and the workflows lack generalization – a workflow that performs well on a specific dataset does not perform well on datasets produced from different studies [37–39]. On the other hand, the SCT gene expression of the same sample, when sample processing procedure and experimental conditions are standardized, are highly reproducible [40,45].

Compared with unsupervised learning, supervised ML classification has superior generalization ability and accuracy. Supervised ML classification systems use algorithms that are logic-based (such as decision trees, rule-based classifiers), network-based (such as artificial neural networks, support vector machines), statistic-based (Bayesian algorithms), or instance-based (such as distance-based or pattern recognition methods) [41]. Supervised ML can perform classification using single-cell gene expression profiles across various studies representing diverse sample processing conditions and experimental settings. Our earlier work demonstrated the potential of artificial neural networks (ANN) to classify healthy PBMC cells in blood samples. In the original study, we achieved the accuracy of 5-class classification (into BC, DC, MC, NK, and TC) of 89.4% [12]. The follow-up study was performed using an improved and expanded data set to perform incremental learning. Several irrelevant data sets were removed, such as DC from non-blood samples (tonsils and tumor ascites) and T cells fixed in methanol, and several new data sets were added to the training set. The 5-class classification accuracy improved to 93% [42]. The introduction of assemblies of ANNs with a new voting function further improved the accuracy of classification to 94.7%, but this required a 100-fold increase of computational processing time.

The previous three studies have demonstrated that high accuracy can be achieved in the 5-class classification of PBMC cell types. The limitation of these studies is that all testing was performed using a single independent (of the training) test set that was annotated by experts. In the current work, we have explored generalization properties of the ANN classification by incremental learning, the effects of data quality on classification accuracy, and have assessed the current accuracy of PBMC classification by ANN. This study is vital for establishing a baseline for comparing healthy samples with those representing various altered conditions, including gene expression changes in disease.

## Methods

This study is an extension of our previous studies [12,42]. The basic ANN classifier is the same as in previous studies. The data sets used for training and testing are different: some of the data sets used in [42] were removed and new data sets were added. Subsequent analysis of data sets used in our previous study indicated that some of the training data represent cells that were processed to the extent that they do not represent healthy PBMC well. The removed data sets include those representing non-malignant cells generated from cancer patients (cutaneous T-cell lymphoma) pre- and post-therapy (GSM3478792 and GSM3558027) [43], *ex vivo* activated of T cells (GSM3430548 and GSM3430549) [44], and cells that represented mixtures of monocytes and dendritic cells (GSM3258347 and GSM3245345). One more high-quality test set, BroadS2, was added to our study (GSE132044, [45]). The first part of the study design included incremental learning with larger and more diverse data sets than in our previous studies [12,42]. The second part of the study involved a comparative validation where all data from one source were used for performance testing while data from other sources were used for training.

### Study design

In the first part of this study, we deployed an incremental learning process for ANN model training and testing as previously described [42]. Five data sets from BroadS1 study were combined to be used as the test set. The training was performed incrementally – data were added to the training set following the order of time of data sets acquisition. Nine cycles of training were done until all training data sets were used. The overall assessment of classification performance was done after Cycle 9. In the final step, we swapped BroadS1 and BroadS2 data sets and assessed the classification accuracy with BroadS2 data set as a test set. The incremental learning process is illustrated in Figure 1.

**Figure 1.**
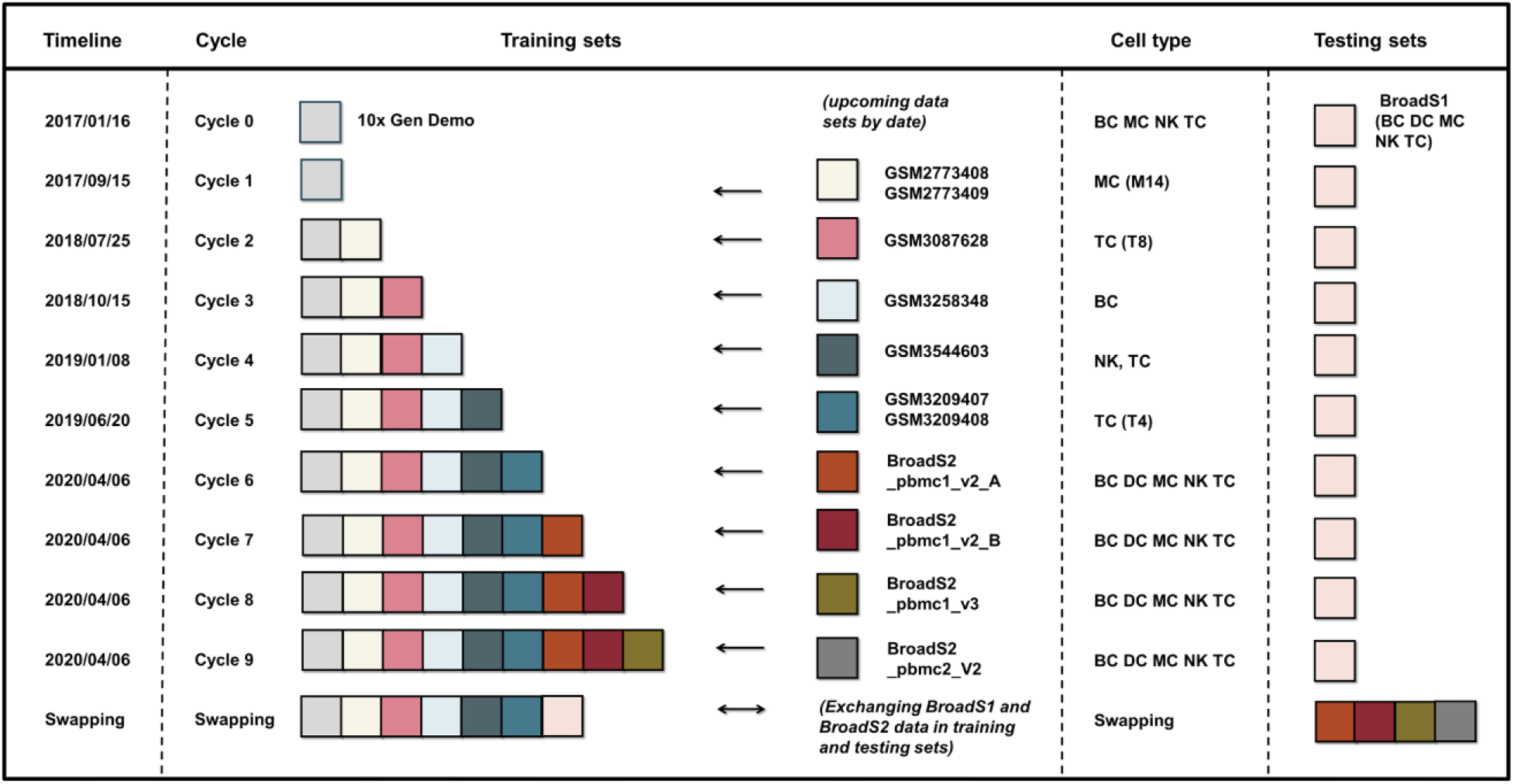
Illustration of the process of incremental learning (training and testing) by adding data sets to the training set and cyclical assessment of classification accuracy. The cycles of learning were ordered by their publication dates to simulate the situation with real life data accumulation. In the final step of incremental learning, BroadS1 and BroadS2 data sets were swapped to observe the reproducibility of ANN results.

Three types of classification tests were performed in each learning cycle (except Cycle 0 and the Swapping Cycle, that do not have upcoming data sets):

- internal 2-fold cross validation on the training set to check the internal consistency of the training data,
- classification accuracy on newly added data sets (upcoming data) before their inclusion in the training set, to check to what extent the gene expression patterns of the added data sets are already represented in the training set,
- classification accuracy of the training set after inclusion of the added data sets using standard independent test set (BroadS1).

The second part of this study involved a comparative analysis of PBMC classification of different training and testing sets. We performed a comparative analysis of the classification of PBMC using four parallel classification models using data sets from our sources:

- Training set: {10x ⋃ GEO ⋃ BroadS2}, testing set: {BroadS1}
- Training set: {10x ⋃ GEO ⋃ BroadS1}, testing set: {BroadS2}
- Training set: {GEO ⋃ BroadS1 ⋃ BroadS2}, testing set: {10x}
- Training set: {10x ⋃ BroadS1 ⋃ BroadS2}, testing set: {GEO}

The comparative analysis involved the assessment of classification accuracy and the interpretation of results using statistical properties of the data sets.

### Data

We selected 58 data sets that represent PBMC from healthy blood samples. These data sets were collected from the NCBI GEO database (www.ncbi.nlm.nih.gov/geo), 10x Genomics demonstration data [27], and Broad Institute Single Cell Portal (singlecell.broadinstitute.org/single_cell). All data sets were processed into our standardized format that has 30,698 features (genes). Most analyses were done using raw data values of standardized features, as provided by the source. Additional validation step was performed with cells from BroadS2 that were subject to quality control: cells that had less than 300 positive features, or less than 670 total counts were excluded, and the results were compared to the results obtained from predictions that used raw data only. 10x demonstration data were generated using standardized 10x scRNA-seq experimental protocol, including validated upstream data analysis [27]. We consider these data sets as reference for PBMC cells processed by PBMC isolation, enrichment (purification), freezing, thawing, and 10x processing.

Thirteen data sets were extracted from the GEO database including GSM2773408, GSM2773409, GSM3087628, GSM3258348, GSM3544603 (seven data sets in this GSM), GSM3209407, and GSM3209408 [21,54–57]. These data sets were generated from PBMC samples extracted from fresh whole blood of healthy donors. These 13 data sets were produced using 10x protocol after PBMC isolation followed by cell sorting by FACS (fluorescent activated cell sorting) or MACS (magnetic activated cell sorting). We obtained two PBMC data sets from Broad Institute Single Cell Portal. The first data set is from the study SCP345, and the second data set is from the study SCP424 (also published in GEO GSE132044 [45]). We named these two data sets BroadS1 and BroadS2, respectively. These data sets were produced using 10x protocol after PBMC isolation followed by annotation of cell types by cell labeling algorithms, and manual labeling correction by experts. These data sets represent a large variety of sample processing, experimental conditions, data analysis approaches, and study purposes. The original test sets (BroadS1) and the newly added set (BroadS2) have multiple repeated SCT measurements of samples from the same individuals at different times, locations, or different chemistry. The same samples processed under the same conditions show high reproducibility. When different chemistry (v2 *vs*. v3 with BroadS2) was used in the 10x protocol, a modest but notable shift in gene expression reproducibility was observed [15]. The summary information on the distribution of cell types across our data sources and their numbers are shown in Table 1. The detailed description of data sets with associated metadata can be found in Supplemental Table 1.

**Table 1.** Summary description of 58 SCT data sets involved in this study. Cell numbers and the number of data sets (values within brackets) are shown per cell type. The data sources are described in the main text. BC – B cells, DC – dendritic cells, MC – monocytes, NK – natural killer cells, TC – T cells. BroadS1 is the original test set.

|  | Cell types - CLASSES |  |  |  |  |  |
| --- | --- | --- | --- | --- | --- | --- |
| SOURCE | BC | DC | MC | NK | TC | TOTAL cells |
| 10x Demo | 10,085 (1) | 0 | 2,612 (1) | 8,385 (1) | 64,341 (6) | 85,423 (9) |
| GEO | 1,760 (1) | 0 | 856 (2) | 309 (1) | 8,789 (9) | 11,714 (13) |
| BroadS1 | 1,660 (1) | 142 (1) | 1,661 (1) | 1,394 (1) | 8,326 (1) | 13,183 (5) |
| BroadS2 | 1,884 (4) | 270 (7) | 2,132 (8) | 842 (4) | 7,164 (8) | 12,292 (31) |
| **TOTAL** | **15,389 (7)** | **412 (8)** | **7,261 (12)** | **10,930 (7)** | **88,623 (24)** | **122,612 (58)** |

The number of data sources in our study is four, the number of data sets is 58. PBMC comprise five main cell types: B cells (BC), dendritic cells (DC), monocytes (MC), natural killer (NK) cells, and T cells (TC). Cell types in our data set have multiple subtypes: NK cells have one subtype; each of BC, DC, and MC has two cell subtypes; TC type has three cell subtypes (Figure 2). TC subtypes are further divided into three sub-subtypes, each for CD8+ T cells and innate-like T cells, and four sub-subtypes of CD4+ T cells. The actual number of PBMC subtypes at multiple levels of ontology is likely to be in hundreds [18]. The total number of cells in our study is 122,612. The test sets have 13,183 cells (BroadS1) or 12,293 cells (BroadS2). The distributions of gene expression values across cells in each data set were visualized. Plotting module pl.violin from SCANPY [46] was used for drawing violin plots.

**Figure 2.**
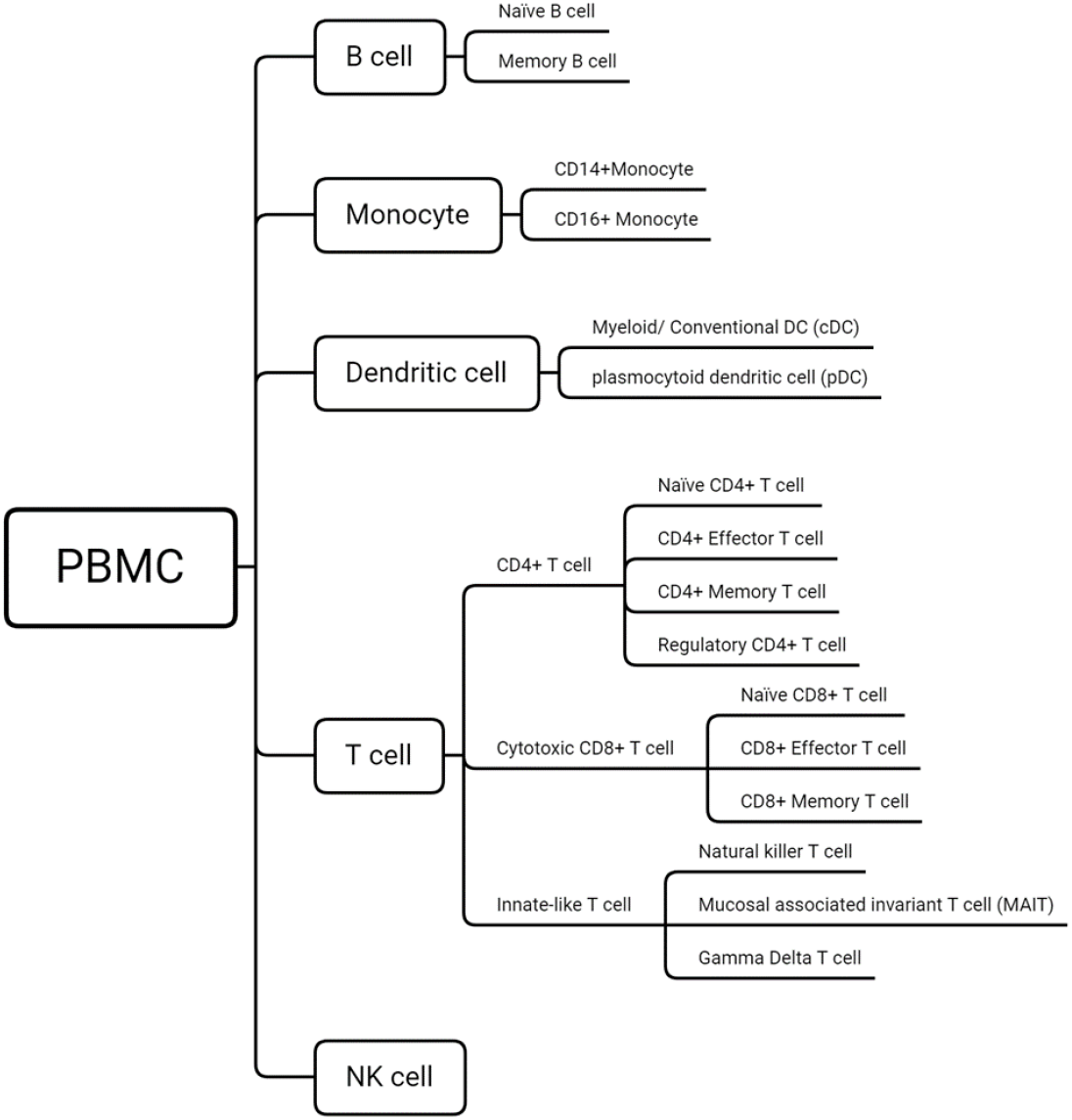
The ontology of cell types and subtypes in our study. The designation of cell subtypes and sub-sub types is provided to show the diversity of cell subtypes used in this study. Because the classification task in this work focuses on the classification of five main types, the descriptions of cell-subtypes and sub-sub types have been omitted.

The data are represented as sparse matrices, where the list of cell identifiers (cell ID) occupies the top row (starts from column 2), and the list of gene names (features) occupies the first column (starts from row 2). The first matrix position (1,1) is blank, while other matrix values represent gene expression counts of a given gene in the given cell determined by the matrix position (gene name, cell ID). Our standardized gene list contains 30,698 genes that are arranged in the same order. Most of the values in an expression matrix are zero.

### Artificial neural networks

For this study, we deployed a fully connected feed-forward artificial neural network (ANN). The ANN system used in this study is illustrated in Figure 3. The multi-layer perceptron classifier MLPClassifier of the scikit-learn [47] python library was used for software implementation. The ANN architecture consists of one input layer, one hidden layer and one output layer (Figure 3 B). The input layer has 30,698 input units corresponding to the 30,698 genes in our standardized SCT data sets (the rows in the sparse matrices). The ten hidden nodes have been chosen to use after exploratory analysis that showed the best balance between the classification accuracy and training speed. The comparison experiments were done using ANN architectures composed of 2, 5, 10, 25, 50 and 100 hidden layer nodes [12]. The output layer is composed of five output units (BC, TC, NK, MC, and DC classes) referring to the respective five PBMC cell types (B cells, T cells, NK cells, monocytes, and dendritic cells). The activation of hidden nodes was done by rectified linear unit (ReLU) function, f(x)=max(0, x). The training data was split into minibatches of size 200. The ANN training used Adam algorithm for first-order gradient-based optimization [48]. The initial learning rate was set to 0.001. In each ANN cycle 90% of the training data was used for training, and 10% were used for validation. The early stopping method was deployed for to prevention of data overfitting. The training stopping condition was when the classification accuracy assessed by validation failed to improve for 11 iterations.

**Figure 3.**
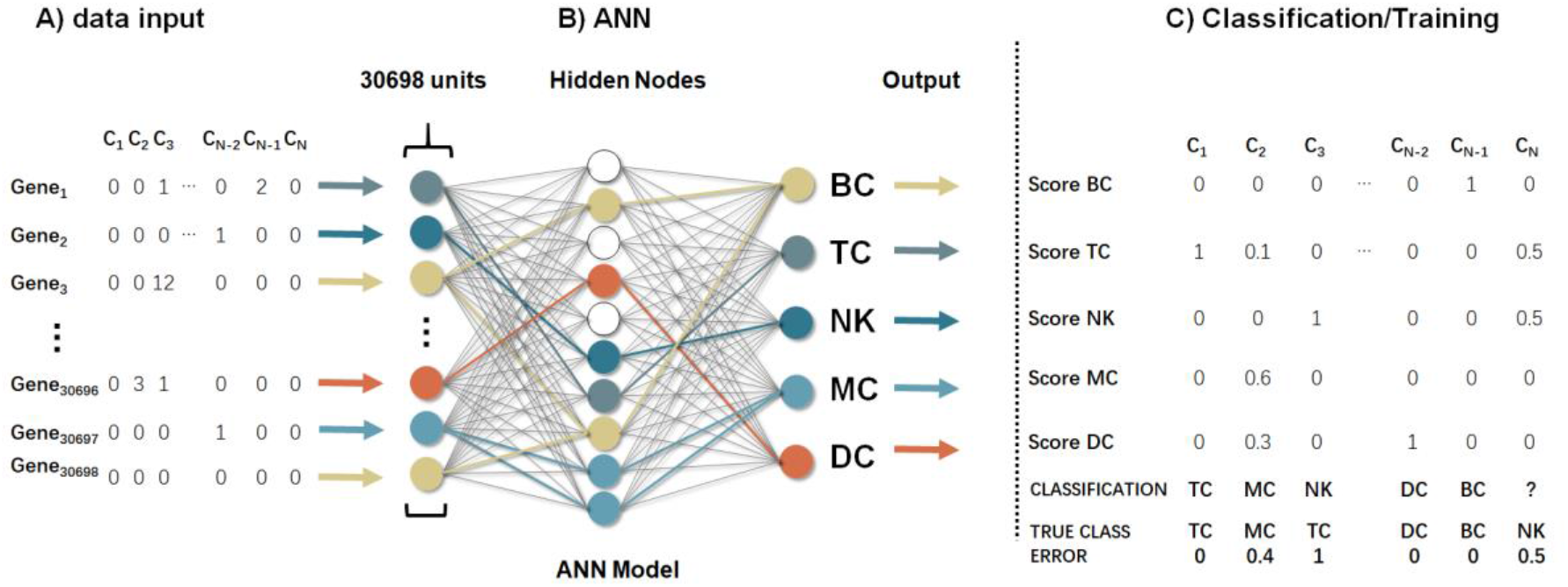
The input data (A), ANN architecture (B), and the output data (C) are shown in this figure. The input data are in the form of sparse matrices where counts are represented by zeroes or positive integers. The architecture is fully connected ANN with 30,698 input units, 10 hidden layer units, and 5 output units, where output units correspond to classes representing major PBMC cell classes. The outputs are represented as matrices of output values that are used in training (by calculating errors) or for prediction of the class of cells of unknown type.

The training data is in the form of large matrices (N × 30,698), where N is the total number of columns – cells in each training step. Gene expression counts of 30,698 genes (Figure 3 A) are in the rows. The output consists of five real numbers obtained from each of the output units, and their sum is V_BC_+V_TC_+V_NK_+V_MC_+V_DC_=1 (Figure 3 C). During training, the weights of the ANN are adjusted and after each adjustment the error is calculated as the sum of the absolute values of the difference between ethe expected value (one for the correct class, and zeroes for incorrect classes) and the actual score of the output units. The ANN training algorithm adjusts the weights between the nodes to minimize the overall output error. For classification, the true class of each cell is unknown, and the predicted class is determined by the maximum value of the five outputs (Figure 3 C).

### Assessment of classification performance

A 5-class confusion matrix was used for the analysis of classifier performance. Confusion matrices were recorded for each cycle of incremental learning. The performance of classification was assessed using measures of Sensitivity/Specificity (Formula 1), Precision/Recall (Formula 2), F1 measure (Formula 3), and the overall Accuracy (Formula 4).

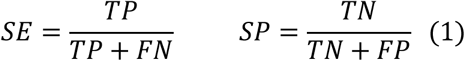

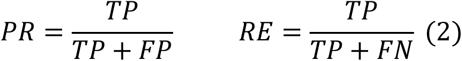

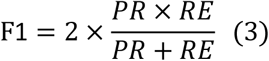

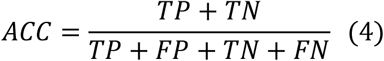

where, SE – sensitivity, SP – specificity, PR – precision, RE – recall, TP – true positives (class positives predicted as positives), TN – true negatives (class negatives predicted as negatives), FN – false negatives (class positives but predicted as negatives), and FP – false positives (class negatives but predicted as positives). SE and RE represent the same entity. Because we performed multi-class classification, accuracy measure was used for the assessment of overall performance, while F1 values were used for the assessment of performance in the classification of individual cell types.

## Results

Density distributions of gene expression within data sets showed a great variety (Figure 4). Data sets from GEO (cells sorted by FACS) show a high median gene expression value (between 3000 and 4000 counts). GEO data sets BC02, MC02, and MC03 showed broad quartile ranges and unimodal density distributions. GEO data sets TC07, TC14, and TC15 showed intermediate quartile ranges with bimodal distributions. On the other hand, GEO data sets NK02, and TC08-TC13 showed high median gene expression values (close to 3000 counts) and narrow quartile ranges, most with bimodal density distributions. NK02 and TC08 data sets showed unimodal distributions and narrow quartile ranges. Bimodal distributions indicate the presence of more than one cell subpopulations.

**Figure 4.**
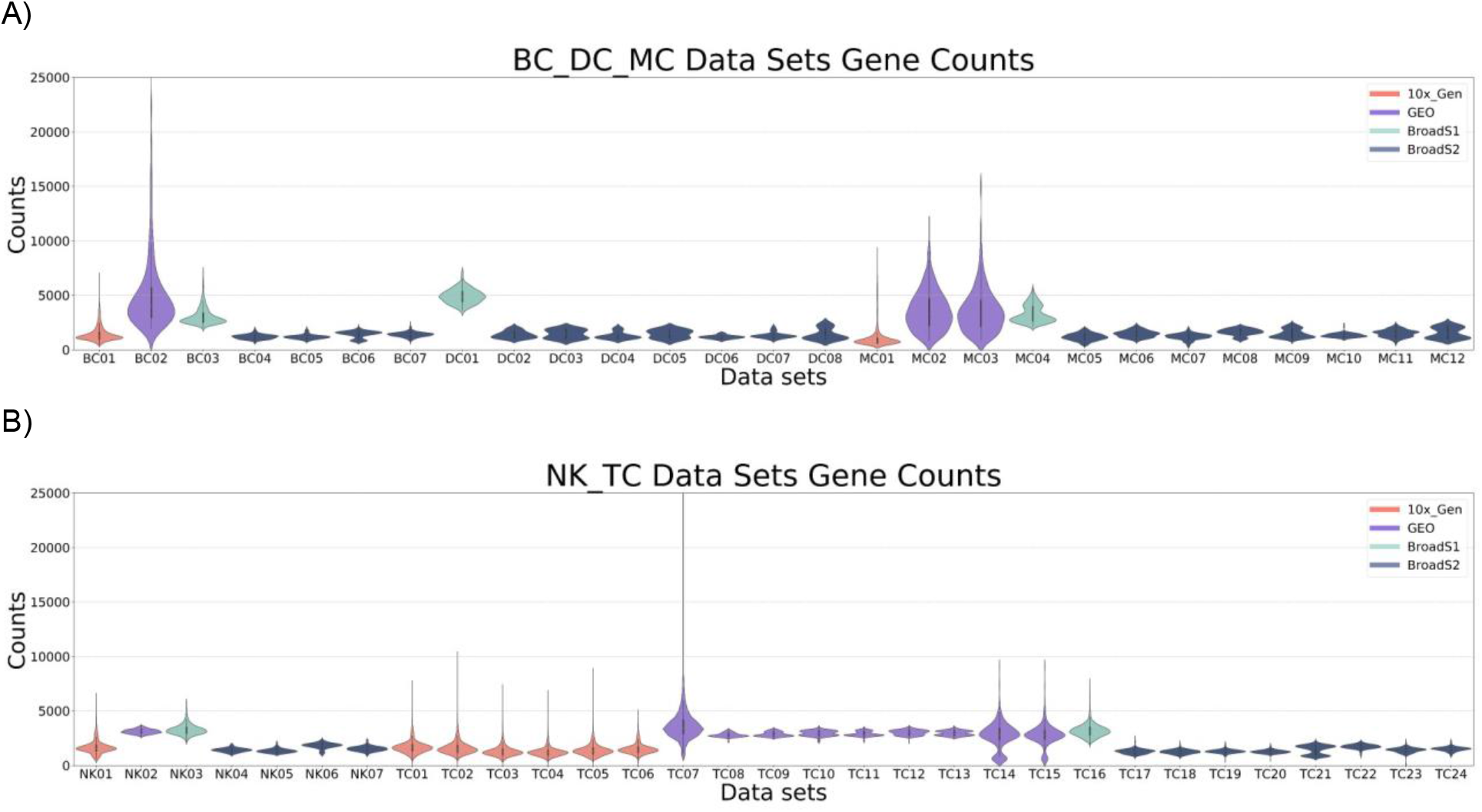
Density distributions of gene expression across 58 data sets used in the current study. A) violin plots of B cells, monocytes, and dendritic cells. B) violin plots of NK cells and T cells. BC01, MC01, NK01, and TC01-TC06 are from 10x demonstration data; BC02, MC02, MC03, NK02, and TC07-TC15 are from GEO data set; BC03, DC01, MC04, NK03, and TC16 are from BroadS1; the remaining data sets BC04-BC07, DC02-DC08, MC05-MC12, NK04-NK07, and TC17-TC24 are from BroadS2. The maximal width of each of the violin plots was set to one (“1”).

Data sets from BroadS1, BC03, DC01, MC04, NK03, and TC16 showed a high median value of gene expression and intermediate breadth of quartile ranges. The majority of BroadS1 cell type data sets showed unimodal distribution, while MC04 showed a bimodal distribution, most likely representing CD14+ and CD16+ monocyte subtypes. We noted that all BroadS1 data have high gene expression counts (4880 ≥ median counts ≥ 2815, across BroadS1 data sets), and high number of positive features (790 ≥ median features ≥ 486) than BroadS2 where expression counts (1843 ≥ median counts ≥1163, across BroadS2 data sets) and positive features (1508 ≥ median features ≥ 611) (Supplemental Table 2). BroadS2 data sets showed wider interquartile ranges than BroadS1 data sets. A large proportion of BroadS2 data sets had shown distinct bimodal distributions (Figure 4 B), indicating that this data may contain distinct subtypes within the indicated cell type. Bimodal counts of gene expression were also observed in T cell data sets from BroadS2 data set and in monocytes from BroadS1.

**Table 2.** The cell type compositions of training and testing sets. The proportions of the main PBMC cell types are shown for healthy range [11,49], training sets, and test sets (BroadS1 and BroadS2).

| CELL TYPE | Healthy Range | Average Training Sets | Test Set BroadS1 | Test Set BroadS2 |
| --- | --- | --- | --- | --- |
| B Cells | 5-15% | 12.08±0.46% | 12.59% | 15.33% |
| Dendritic Cells | 1-2% | 0.07±0.09% | 1.08% | 2.20% |
| Monocytes | 10-30% | 4.10±0.63% | 12.60% | 17.34% |
| NK Cells | 5-10% | 9.09±0.36% | 10.58% | 6.85% |
| T Cells | 40-70% | 74.67±0.76% | 63.15% | 58.28% |

### Incremental learning

The average composition of the training sets and the compositions of test sets are shown in Table 2. The composition of the training sets is stable across cycles (Fig. 5). Test sets match well the healthy ranges [11,49] while DC were severely underrepresented in the training sets, monocytes were underrepresented, and T cells were overrepresented (Table 2). The DC were included in the training set only in Cycles 6-9 and their representation in the training set remained low, approximately 10- to 20-fold lower than their representation in test sets. The training set in Cycle 0 included only samples that were from 10x demonstration data – processing of these cells included PBMC extraction, purification by bead-enrichment, freezing, thawing, and 10x processing. Cycles 1-6 included the addition of cells sorted by FACS or MACS to the 10x data set. Testing in all cycles was performed using minimally processed (PBMC extraction and freezing) data set BroadS1. The final round, swapping, involved two steps: a) training data set included 10x, GEO and BroadS2 data, and testing was done using BroadS1 and b) training set included 10x, GEO, and BroadS1 data, and the entire BroadS2 data set was used for testing.

**Figure 5.**
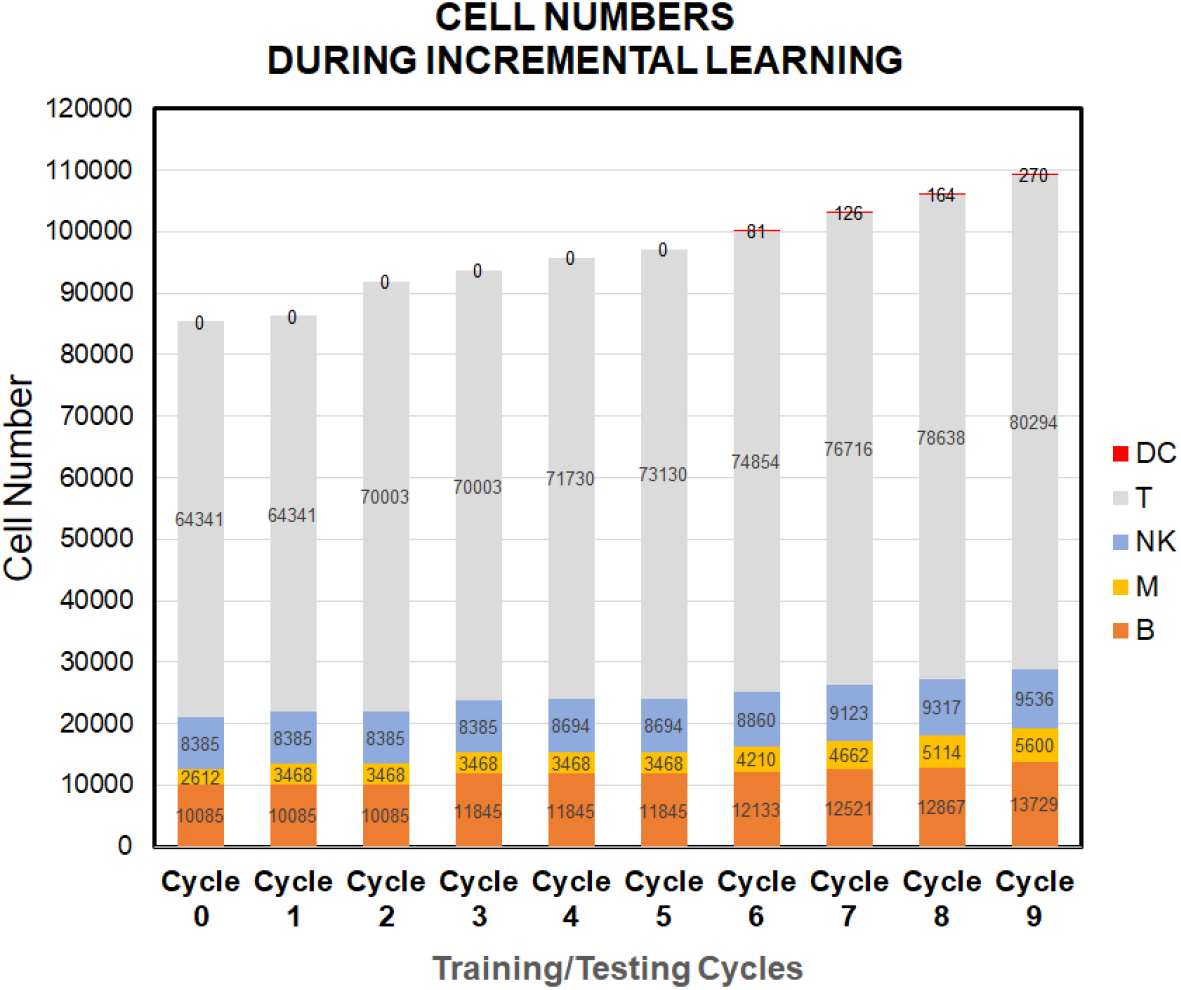
Data sets used in training cycles appear in the time sequence as we acquired them. The increase in the number of cells in training sets was gradual and the proportions of cell types were stable. The new sets of cells tested in the current cycle were appended to the subsequent training set. For example, monocytes from GSM2773408 (333 cells) and GSM2773409 (235 cells) were classified using the training set from 10x dataset (cycle 0), then were included in the training set for Cycle 1.

The results of ANN classification results are shown in Fig 6. The internal cross-validation showed reproducibly high accuracy ranging from 99.9% to 99.2%. The accuracy of classification of new independent data sets was initially low (66.4% in cycle 0 and 32.1% in Cycle 1, then it rapidly increased and stabilized between 90% and 99% from Cycle 4. The external validation with BroadS1 data set showed low accuracy of classification in Cycles 0 and 1, followed by a rapid increase to 87.6%, followed by a gradual improvement in accuracy that reached 94.3% in Cycle 9. The swapping step, where BroadS2 was used as a test set showed the accuracy of internal cross-validation of 98.9% and external validation accuracy of 92.8%. Taken together, the results indicate that the overall accuracy of 5-class classification is between 93 and 94%.

**Figure 6.**
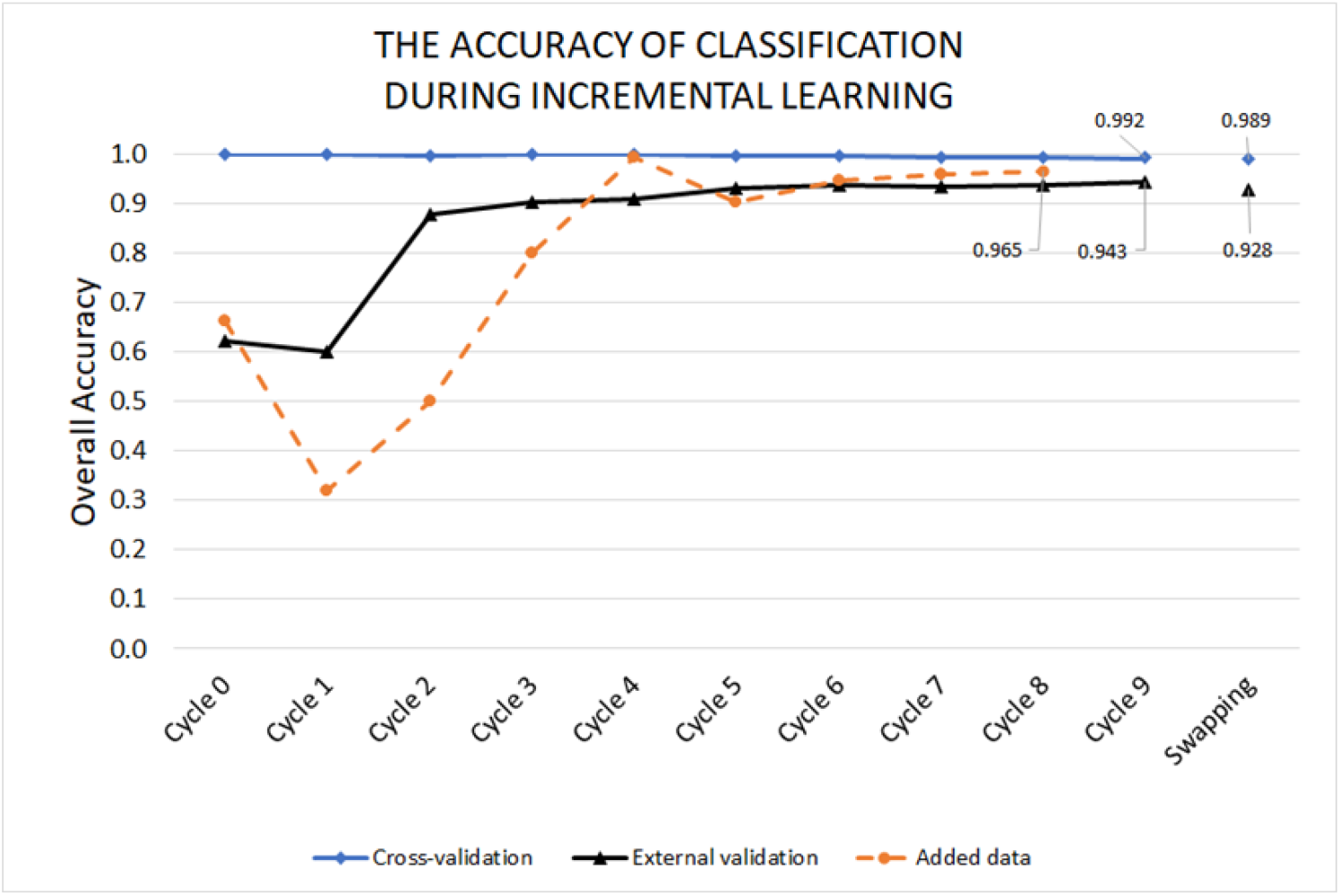
The internal cross-validation showed extremely high accuracy (≥98.9% in all cycles). After early instability (Cycle 1) the classifier starts converging towards internal cross-validation line. With the increase of the number of data sets added to the training set, new data files are predicted with steadily increasing accuracy (added data line). The swapping step showed that the overall accuracy of the current system is within the range of 93-94%.

The Cycle 9 and the swapping produced results for EXP 1 and EXP 2 (Figure 6 and Table 3). The remaining part of our study involved training of the ANN classifier by 10x+BroadS1+BroadS2 and testing with GEO data (EXP 3, Table 3), and training of the ANN classifier by GEO+BroadS1+BroadS2 and testing with 10x data (EXP 4, Table 3). Sample processing alone has a profound effect on gene expression (Table 3). The prediction model trained on combination of minimally processed samples combined with samples processed by enrichment or cell sorting can be used for high accuracy prediction of minimally processed samples.

**Table 3.** Classification accuracy for modeling experiments where the testing set derived entirely from one source, while training sets were combined from other sources. The results also show F1 measure for individual cell types. Further details are available in Supplemental Tables 4 and 5.

| EXP | TEST SET* | SAMPLE PROCESSING | CLASSIFICATION ACCURACY | F1 VALUES |
| --- | --- | --- | --- | --- |
| 1 | BroadS1 | Separation | 94.3% | BC – 0.956, DC – 0.807<br>MC – 0.975, NK – 0.786<br>TC – 0.964 |
| 2 | BroadS2 | Separation | 92.8% | BC – 0.956, DC – 0.022<br>MC – 0.973, NK – 0.739<br>TC – 0.949 |
| 3 | 10x | Separation, Enrichment | 46.3% | BC – 0.721, DC – 0.000<br>MC – 0.009, NK – 0.253<br>TC – 0.591 |
| 4 | GEO | Separation, FACS or MACS | 75.7% | BC – 0.773, DC – 0.000<br>MC – 0.805, NK – 0.236<br>TC – 0.835 |
| 5 | BroadS1 | Separation | 94.0% | BC – 0.961, DC – 0.230<br>MC – 0.964, NK – 0.784<br>TC – 0.965 |
| 6 | BroadS2 | Separation | 88.1% | BC – 0.876, DC – 0.000<br>MC – 0.971, NK – 0.592<br>TC – 0.927 |

\*EXP 1-4 involve three training sets and one testing set. EXP 5 and 6 involve only BroadS1 and BroadS2 data sets.

The overall performance of classification differs between individual cell types (Table 3, EXP 1 and 2): B cells, monocytes, and T cells show high accuracy with F1 values exceeding 0.95. Classification performance of NK cells shows lower accuracy with F1 value in the vicinity of 0.75. Quality control of BroadS2 data set (removal of cells that have total counts lower than 670 or number of positive features lower than 300) did not affect classification performance (EXP 2a and EXP 2b, Supplemental Table 5). Classification of dendritic cells was unstable, F1=0.81 in EXP 1 and 0.02 in EXP 2 (Table 3). When two-fold external validation with BroadS1 and BroadS2 data sets were performed (EXP 5 and 6, respectively), the overall accuracy in EXP 5 was 94%, and in EXP 6 was 88.1%. The inclusion of data sets with high median gene expression (GEO, 2700-3500 and BroadS1, 2800-4900, Supplemental Table 2) in the training data set results in lower cell classification accuracy (EXP 2 as compared to EXP 1, EXP 3 as compared to EXP 4, and EXP 6 as compared to EXP 5). The lack of data produced by separation + enrichment in the training set in EXP 3 coincides with very low classification accuracy (46.3%, Table 3). Similarly, the lack of data produced by separation + FACS/MACS in the training set in EXP 4 coincides with low classification accuracy (75.7%, Table 3). On the other hand, adding BroadS1 to the training set in the swapping step, as compared to Cycle 5, further improved the classification accuracy tested on BroadS2 (91.6-92.8%, EXP 7 and EXP 2, Supplemental Tables 3, 4 and 5).

## Conclusions

Overall, our results demonstrate that supervised ML is a viable option for classifying cell types from single cell expression data. Patterns that are characteristic for cell types are preserved in single cell gene expression data even when the single cell samples are processed using different processing steps. Data sets derived from minimally processed samples (PBMC separation only) alone cannot be used to predict cell type from samples that are additionally processed (we achieved a prediction accuracy of 46.3% for enriched and 75.7% for sorted cells, Table 3). However, gene expression patterns characteristic of a given cell type are preserved in samples that have additional processing steps and these sets can be used for accurate predictions of minimally processed samples (93% accuracy on BroadS1 data set was achieved by training set consisting of 10x and GEO data, Figure 6 and Supplemental Table 3). For broad application. The training data set – the reference set – is composed of multiple data sets that represent various sample processing conditions and contain sufficient biological variability. The ANN classifier is robust – the system can tolerate a proportion of cells that have gene expression lower than quality control thresholds (in our studies it is 670 for gene expression counts and 300 for positive features).

Two-fold internal cross-validation has shown that once a data set is added to the training set, the patterns contained in that set will be remembered by the classifier. The classifier generalizes well, and generalization properties improve with the addition of new data. Once a data set representing a particular cell type and sample processing protocol is added to the training data set, the ANN will learn this data type. When data sets where a particular cell type, biological condition, and experimental processing protocol is well, that is very high (≥98.9%, Fig. 3).

The overall classification performances in EXP 1 and 2 (Table 3) are satisfactory (94.3% and 92.8%), but in EXP 3 and 4 are not (46.3% and 75.7%). Training data used in EXP 1 and EXP 2 are representative of all three sample processing protocols: i. separation (of PBMC), ii. separation + enrichment, and iii. separation + cell sorting. Training data used in EXP 3 did exclude separation + enrichment protocol data that was used for generating test data in the same experiment. Similarly, test data in EXP 4 were generated using separation + cell sorting protocol, while the corresponding training data represented samples produced by other processing protocols. A well-established classification theory concept in ML is that the training set must be representative of the variability that is present in real cases. Our results clearly show the effects of the training sets that are not fully representative. A problem for SCT is that processing steps such as enrichment or cell sorting are part of the experimental validation of results that are missing in minimally processed samples. Our results of EXP 1 and 2 show high accuracy of classification but cannot be validated directly by experiments. On the other hand, the cell type in EXP 3 and 4 is known, but the classification accuracy is low if similar data are not present in the training set.

EXP 1-8 (Supplemental Table 5) results indicate that the average gene expression level of data sets used in training has an influence on classification accuracy, particularly in situations where the training set is limited. The results indicate that the classification of cell types is better in data sets that have moderate gene expression levels, with gene counts between 1000 and 2000 per cell. This observation needs further study to confirm the actual influence of gene counts on classification accuracy. The analysis of factors that possibly influence prediction accuracy in this study is presented in the Discussion section.

In summary, we have demonstrated that ANN, a supervised ML method, is capable of high accuracy classification of five main cell types of healthy PBMC. The accuracy is very high for B cells, Monocytes, and T cells. The classification accuracy of NK cells is lower, because of their similarity with subsets of T cells (such as NK-like T cells, subsets of CD8+ T cells, and subsets of innate T cells). This problem was noted in [27], where the authors reported that it was challenging to separate cytotoxic T cells and NK cells since they have overlapping feature spaces. The accuracy of the classification of DC is low because of the underrepresentation of DC in the training sets, and this problem should be overcome by adding additional DC samples.

## Discussion

This work demonstrates the potential of supervised ML methods to classify single cells from their gene expression counts. We achieved the 5-class classification accuracy of 94% using 58 data sets derived from healthy PBMC that were processed by different experimental procedures applied to PBMC samples. An important finding is that once a dataset representative of a cell type, condition (in this case healthy PBMC), and a specific sample processing protocol is added to the training set, similar data sets will be classified with very high accuracy (>99%).

Several factors limit the accuracy of our 5-class classification of PBMC. They include lack of training data (for DC) and similarity of cell subtypes with cells from other classes (NK cells), and training data with high median gene counts. Additional factors include undefined classes or subclasses of cells that are normally found in peripheral blood but are not included in current training set. Such cells, for example, include CD34+ cells (circulating hematopoietic cells that may represent between 0.1 and 0.3% of PBMC [49]. Natural killer T (NKT) cells have markers of both T cells (CD3+) and NK cells (CD56+) and are present in circulating PBMC [50] and can easily be confused with NK cells. On the other hand, CD8+ NK cells [51] share properties with cytotoxic T cells. Given the similarity of gene expression profiles, is not surprising that in our study, 3.6% (297) of T cells from BroadS1 and 4.8% (345) of T cells from BroadS2 were classified as NK cells. Conversely, 18.2% (254) of NK cells from BroadS1 and 17.2% (145) of NK cells from BroadS2 were classified as T cells. FACS sorting showed that NK cells from 10x data were 92% pure, while CD8+ cytotoxic cells were 98% pure. Further investigation, including advanced clustering methods [such as 52,53] and the analysis of misclassifications, will be pursued to improve PBMC classification.

One challenge for the classification of cells from SCT data arises from the need for experimental validation of cell types as opposed to expert annotation in minimally processed samples. Experimental sample processing steps such as bead enrichment (negative selection) produce homogeneous samples (one cell type or subtype) whose purity can be verified by cell sorting. Alternatively, cells can be sorted by FACS or MACS procedures that help sort cells, and provide a measurement of purity, percentage of contaminating cells, and cell properties [*eg.* 54]. Depending on the purpose of single cell study, various sample processing workflows may be deployed (Fig. 7). The difficulty with processed samples is that each processing step induces changes in gene expression profiles. These profile changes are significant, and they prevent direct comparison of cells from studies that follow different protocols. Minimally processed samples have similar gene expression to the native blood cells. The annotation of single cells in this case, is done by various tools that utilize gene expression markers and are normally inspected and corrected manually, introducing annotation bias. Protein markers and gene expression markers do not match perfectly, the expression of proteins and corresponding mRNA significantly correlate only in about one-third of targets [59,60]. Since SCT data sets are sparse and a large proportion of expressed genes are missing, simple marker-based assignments are insufficient. A selection of *in silico* methods is needed in combination with experimental validation for conclusive assignment of cell types and subtypes.

**Figure 7.**
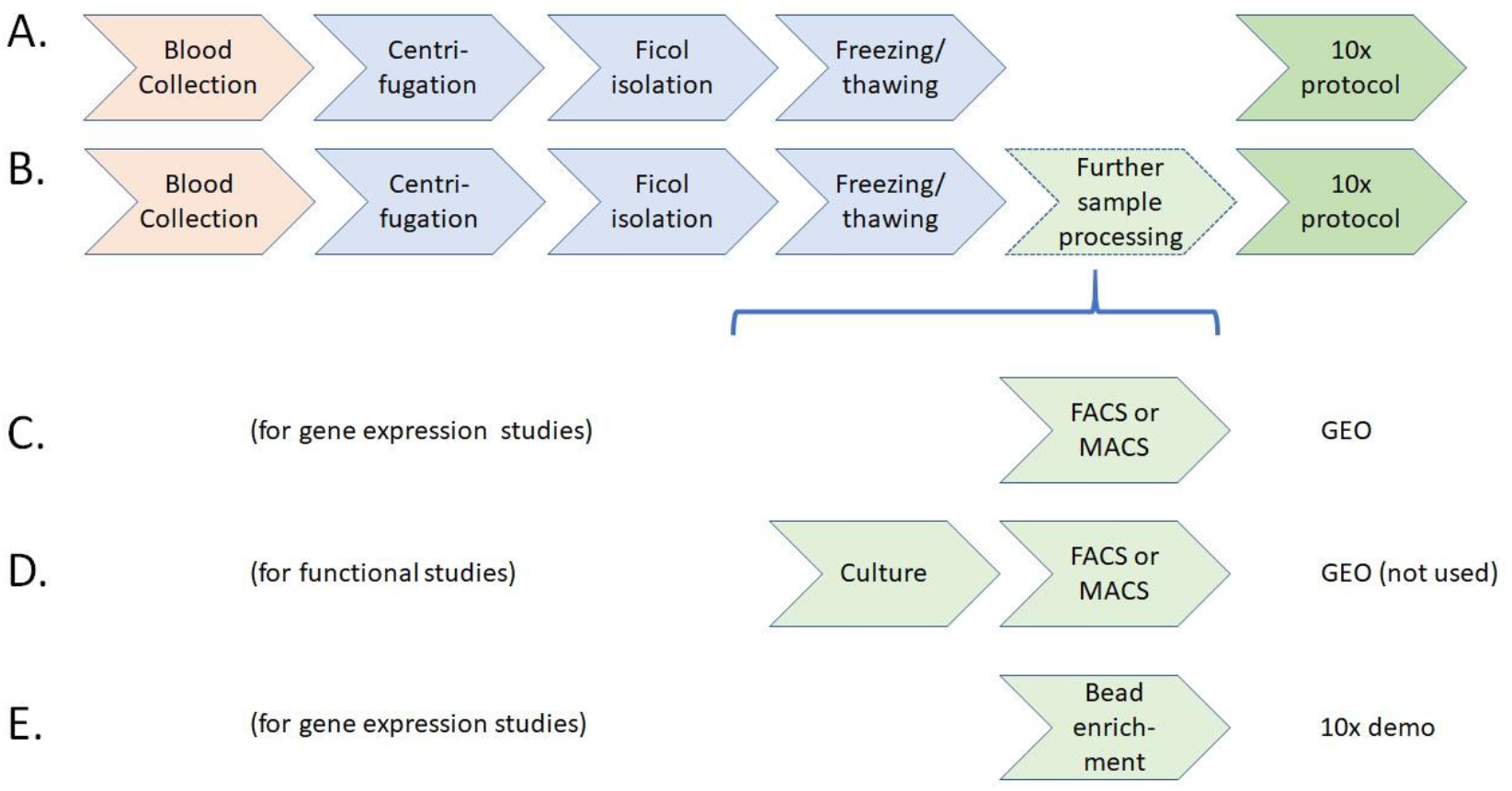
Sample workflows relevant for our study: A. Workflow involving minimally processed samples (BroadS1, BroadS2), B. Generic flow for 10x studies, C. Workflow of samples processed by FACS or MACS, may include multi-step processing (GEO data sets in our study), D. Workflow for functional studies, PBMC samples are often cultured overnight along with bioactive agents, followed by FACS/MACS, E. Workflow using negative selection by bead enrichment (used in 10x demonstration study). Workflow D was not used in this study because culturing with bioactive agents generates cellular responses not relevant for profiling of PBMC from healthy blood.

Supervised ML has distinct advantages in comparison with unsupervised clustering when used for classification tasks. The main advantage is that once reference sets are available, standardized analysis can be performed across samples that represent various biological conditions. Single cell technologies applied to PBMC require the ability to analyze minimally processed samples directly and accurately and reproducibly determine cell types, subtypes, and their status from single cell expression profiles. To achieve this goal, we need standardized sample processing workflows and SOP of upstream single cell analysis and supervised ML methods for downstream analysis. Several sample processing protocols were demonstrated as reproducible and are available (see support.10xgenomics.com/single-cell-gene-expression/sample-prep). SCT samples can be analyzed using existing SOPs and they yield highly reproducible results (as demonstrated, for example, in [15,45]).

Given that the SCT part is stable, supervised ML requires that training data are representative of all major sample processing protocols. Supervised ML analysis can classify any future sample collected, prepared, and analyzed using one of the validated protocols with the expected accuracy. Our results indicate that the accuracy of classification from validated protocols should be above 98%, which matches cell purity from standard cell sorting methods. New sample processing protocols can be validated by splitting minimally processed samples, perform supervised method (such as ANN) classification on one partition of the sample, and performing additional processing steps to confirm the numbers or proportions of cell types in the second partition. In this study we have defined a reference data set for 10x PBMC 5-class classification that provides 94% accurate classification. Our future goal is to refine classification of DC, by increasing the number of DC data in the training set and resolve ambiguities between NK cells and subsets of T cells (non-classical T cells and CD8+ T cells) that are misclassified due to their gene expression similarity.

## Supporting information

Supplemental Table 1. Metadata describing samples as described by the sources.

Supplemental Table 2. The results of basic statistical analysis of the data sets.

Supplemental Table 3.The assessment of classification performance for incremental learning by cycles and steps.

Supplemental Table 4. Confusion matrices for incremental learning by cycles and steps.

Supplemental Table 5. The assessment of classification performance for specific simulations EXP 1 through EXP 8.

## Abbreviations

ANN: artificial neural network
BC: B cells
DC: dendritic cells
FACS: fluorescence-activated cell sorting
MACS: magnetic-activated cell sorting
MC: monocytes
ML: machine learning
NK: natural killer cells
PBMC: peripheral blood mononuclear cells
scRNAseq: single-cell RNA sequencing
SCT: single-cell transcriptome
SOP: standard operating procedures
TC: T cells

## Acknowledgements

Jiahui Zhong is supported by an UNNC IAMET scholarship.

## Authors’ contributions

VB, ZC, and DK conceived the idea. VB, LTC and GZ designed the study. JZ, ML, and VB performed the analysis. VB, JZ, HJ, DK and ML discussed the results. ML, VB, LTC and GZ created datasets and Web Page for data sharing. All authors contributed to writing of the paper. JZ and VB took the lead in writing the manuscript.

## Availability of data and materials

All datasets from this study are available at http://projects.met-hilab.org/SCTdata/PBMC001

## Ethics approval and consent to participate

Not ethic approval was required for this study.

## Supplementary information

Supplemental Table 1. Metadata describing samples as described by the sources.

Supplemental Table 2. The results of basic statistical analysis of the data sets.

Supplemental Table 3. The assessment of classification performance for incremental learning by cycles and steps.

Supplemental Table 4. Confusion matrices for incremental learning by cycles and steps.

